# Proteoform Identification by Combining RNA-Seq and Top-down Mass Spectrometry

**DOI:** 10.1101/2020.05.27.119644

**Authors:** Wenrong Chen, Xiaowen Liu

## Abstract

In proteogenomic studies, genomic and transcriptomic variants are incorporated into customized protein databases for the identification of proteoforms, especially proteoforms with sample-specific variants. Most proteogenomic research has been focused on combining genomic or transcriptomic data with bottom-up mass spectrometry data. In the last decade, top-down mass spectrometry has attracted increasing attention because of its capacity to identify various proteoforms with alterations. However, top-down proteogenomics, in which genomic or transcriptomic data are combined with top-down mass spectrometry data, has not been widely adopted, and there still lack of software tools for top-down proteogenomic data analysis. In this paper, we introduce TopPG, a proteogenomic tool for identifying proteoforms with genetic alterations and alternative splicing events. Experiments on top-down proteogenomic data of DLD-1 colorectal cancer cells showed that TopPG can confidently identify proteoforms with sample-specific alterations.

## 1. Introduction

Top-down mass spectrometry (MS) has been widely used in proteoform identification and characterization because of its ability to sequence whole proteoforms^1^. In a top-down MS experiment^2^, intact proteoforms are separated by a protein separation platform such as a liquid chromatography (LC) system, and then analyzed by tandem mass spectrometry (MS/MS) to generate MS1 spectra of proteoforms and MS/MS spectra of proteoform fragments. Each of the mass spectra contains a list of peaks measuring the mass-to-charge ratios (*m/z* values) and abundances of proteoforms or their fragments ^3,4^.

Database search is the dominant method for top-down MS-based proteoform identification and characterization ^5,6^. In this method, a query top-down MS/MS spectrum is searched against a protein sequence database to identify a proteoform that best explains the peaks in the spectrum. Proteoforms identified by top-down MS are often modified forms of protein sequences in the database. These proteoforms can be further characterized to localize their alterations and find their combinatorial alteration patterns using probabilistic models ^7, 8^.

A main challenge in proteoform identification is that proteoforms contain various alterations, such as sequence mutations, splicing events, and post-translational modifications (PTMs) ^9, 10^. Intact proteoforms with hundreds of amino acids tend to have more alterations than short peptides analyzed in bottom-up MS. However, most protein databases used in proteomics studies, such as UniProt proteome databases, contain only reference protein sequences, not proteoforms with various alterations ^11^.

Many computational methods have been proposed for identifying proteoforms with alterations, which can be divided into three categories ^12^. In the first approach, sequence annotations in protein knowledge bases or genomic or transcriptomic data are used to build an extended database that includes proteoforms with alterations ^13^. A spectrum of a proteoform *A* with alterations can be easily identified if the extended database includes the sequence of *A*. Even if the sequence is not included, but another proteoform *B* that is similar to *A* is included, the extended database also facilitates the identification of the spectrum because proteoform *B* is a good reference sequence.

In the second approach, an open search method is used to identify proteoforms with unexpected alterations, in which the mass shifts of the alterations are reported. This method has been widely used in bottom-up ^14^ and top-down MS ^13, 15^ data analysis. It is capable of identifying proteoforms with one unexpected alteration by using unmodified proteoform fragments. The main limitation of the method is that only one unexpected mass shift can be identified in proteoforms.

In the third approach, spectral alignment algorithms are used to align mass spectra against protein sequences ^16, 17^, which are capable of identifying proteoforms with several unexpected alterations. Some alignment algorithms allow users to provide several expected PTMs ^7, 15, 18, 19^, making it feasible to identify highly modified proteoforms, such as histone proteoforms.

Most existing tools adopt one or two of the above approaches. ProSightPC ^13^ uses both the extended database and open database search methods; TopPIC ^20^, TopMG ^21^, SPECTRUM ^22^ and MS-PathFinder ^15^ use the open search and spectral alignment methods ^23^. Because of the complementary strengths of the approaches, combining extended databases and the other two approaches can increase the sensitivity in the identification of complex proteoforms.

Extended proteoform databases are built using protein annotations and/or proteogenomic methods. Building proteoform databases using annotations in protein knowledge bases, such as UniProt ^11^ and neXtprot ^24^, has several limitations. First, we are still at the early stage of studying proteoforms and lack complete annotation of proteins ^7^, making it impossible to build a comprehensive extended database using protein annotations. For example, less than 1% sequences in the Swiss-Prot database contain annotated PTMs ^25^. Second, annotated alterations in these knowledge bases are not sample-specific. Many annotated alterations exist in some cell types or conditions, but not the sample being studied.

Proteogenomic methods utilize genome annotations and DNA/RNA sequencing data to build extended proteoform databases ^26^. While proteoform databases built with genome annotations only are not sample specific, sample specific proteoform sequences can be incorporated into extended database when genome sequencing or RNA-Seq data are available. For example, based on RNA-Seq data, a customized proteoform database can be built to include sequences with sample-specific genetic alterations and alternative splicing events ^27, 28^. Using such a database in MS data analysis increases proteoform identifications with genetic alterations and alternative splicing events.

Proteogenomic methods also facilitate proteoform characterization in top-down spectral interpretation. Although top-down MS is capable of identifying complex proteoforms, most modified proteoforms are not characterized because top-down MS/MS spectra often lack enough fragment ions. Genomic and transcriptomic data provide additional information for identifying and characterizing alterations in proteoforms. If a mutation in a protein sequence is supported by both MS and transcriptomics data, the confidence of the identification is significantly increased ^27^.

Many proteogenomic pipelines and software tools have been proposed for combining genomic or transcriptomic data with bottom-up MS data ^29, 30^. Most tools like customProDB^31^, MutationDB, MSProGene^32^, PGA^28^, and and JUMPg ^33^ generate sample specific databases including sequences with mutations and/or splice junctions from RNA-Seq data. SpliceDB ^34^ and Splicify ^35^ generate customized protein database with only splicing variants. SpectroGene ^36^ builds protein sequence database by six-frame translation from open reading frames in genomes. PGTools ^37^ utilizes annotations of Ensembl ^38^ to build various protein sequence databases with mutations and splicing events. Askenazi et al. developed a proteogenomic tool called PGx, which maps identified peptides onto their putative genomic coordinates ^39^.

Only several studies have been carried out in top-down proteogenomics despite its potential to identify complex proteoforms with sample specific alterations. When the genomic annotation of the organism being studied is unavailable or incomplete, proteoform databases are generated from genomic data of prokaryotic organisms using six frame translation ^36^ or from transcriptomic data using *de novo* assembly ^40^. When genomic annotation is available, the annotation is utilized to align RNA-Seq reads to the reference genome to identify genomic alterations and novel splicing junctions ^40, 41^. One pilot study of patient-derived mouse xenograft samples of human breast cancer identified 41 single amino acid variations and 11 novel splicing junctions using a top-down proteogenomic approach ^41^.

In this paper, we propose TopPG, a software pipeline for combining RNA-Seq and top-down MS data for proteoform identification. TopPG builds sample-specific proteoform sequence databases using genomic alterations and splicing junctions identified by aligning RNA-Seq reads to the reference genome. We assessed TopPG on a top-down MS data set of DLD-1 colorectal cancer cells and demonstrated that TopPG increased the number of proteoform identifications and identified many proteoforms with sample-specific mutations and splicing events compared with reference databases from ENSEMBL ^38^.

## 2. Methods

### 2.1 Data sets

An RNA-seq data set of DLD-1 colorectal cancer cells (SRR6929326) was download from the sequence read archive (SRA), which contain 48.8 million paired short reads (150 bp). The RNA sample was prepared using the Illumina TruSeq RNA Sample Preparation protocol and deeply sequenced with an Illumina HiSeq 3000 ^42^. Phred quality scores (Q score) reported by FastQC (version 0.11.8) ^43^ showed that more than 90% of the short reads reached Q30 (99.9% base call accuracy).

A top-down MS data set of DLD-1 cells was downloaded from the MassIVE repository (ftp://massive.ucsd.edu/MSV000079978/) ^44^. In the MS experiment, protein sample of DLD-1 cells was first separated into 12 fractions by a gel-eluted liquid-fraction entrapment electrophoresis (GELFrEE) fractionation system. Then the first 8 fractions were analyzed by reverse-phase liquid chromatography coupled with a 21 T FT-ICR mass spectrometer. A total of 14588 MS/MS spectra were collected. Details of the experiment can be found in ref. 44.

### 2.2 A pipeline for building proteoform sequence databases with genomic alterations and alternative splicing events

We developed a top-down proteogenomic pipeline that uses RNA-Seq data to build customized proteoform sequence databases. In the pipeline, RNA short reads are aligned to a reference genome to identify genomic alterations and alternative splicing events, based on which two proteoform sequence databases are generated: one containing proteoforms with genomic alterations, and the other proteoforms with alternative splicing events.

#### 2.2.1 RNA-seq data analysis

The GATK pipeline (version 4.1.0.0) ^45^ is used for the alignment of RNA-seq short reads. In the alignment process, short reads are aligned to the hg38 reference genome (downloaded from the GATK resource bundle) using the two-pass mode of STAR (version 2.7.0.c) ^46^, in which splicing junctions identified in the first round are used to guide the second round of alignment. Picard (version 2.18.26) ^47^ is used to remove duplicates from the short read alignment BAM file, then sort and add indexes to the file. Finally, we use SplitNCigarReads from GATK to remove skipped regions in sequence alignment, and use the BQSR method to recalibrate base quality scores of the short reads. The parameter settings and commands of the tools are given the supplementary material.

The HaploTypeCaller tool in GATK is used to call variants from short read alignment files. The minimum phred-scaled confidence threshold is set to 20 (the error rate is smaller than 1%); soft-clipped bases are excluded to reduce false positives. To further optimize the sensitivity and specificity, VariantFiltration is employed to filter out single nucleotide variant (SNV) clusters with 3 SNVs in a window of 35 bases ^48^.

#### 2.2.2 Building protein sequence databases with genomic variants

All SNVs, insertions and deletions (indels) reported by HaploTyperCaller are annotated by ANNOVAR ^49^ (version April 16, 2018). A protein sequence often contains multiple genomic variants, and many similar proteoform sequences with variants can be generated from the protein sequence. To address the problem, we follow the method proposed by Kolmogorov et al. ^36^ to generate *short protein sequence segments* with variants. Because most mass spectrometers can identify only proteoforms with a molecular mass less than 50 k Dalton (Da), we choose 600 as the length of short protein sequence segments. Given a transcript with *n* nucleotides, we split the transcript sequence into overlapping segments *S*_1_, *S*_2_, …, *S_k_* with a window of length *L*=1800, where 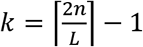. The overlapping region of two segments *S_i_* and *S_i+1_*, for 1 ≤ *i* ≤ *k* − 1, is 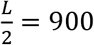. Each segment with 1800 nucleotide bases is translated to a protein segment with 600 amino acids.

Genomic variants (SNVs and Indels) reported by ANNOVAR are divided into two types: homozygous and heterozygous. For a transcript *T* with homozygous and heterozygous variants, we generate a transcript *T*’ with all the homozygous variants, and then split *T*’ into overlapping segments *S*_1_, *S*_2_, …, *S_k_* with length *L*. If *S_i_* (1 ≤ *i* ≤ *k*) contains *j* heterozygous variants, we generate *j* transcript segments from *S_i_*, each of which contains one heterozygous variant. Note that a single position in the human genome may have more than one heterozygous SNV and that the number of heterozygous SNVs may be larger than the number of SNV sites. Finally, we obtain a protein segment from each generated transcript segment if it contains genomic variants. If a segment *S_i_* (1 ≤ *i* ≤ *k*) does not contain any variants, it is not used to obtain a protein segment. When a transcript segment has frameshift indels, we find the first downstream stop codon with the new frame and use the new frame and the stop codon to generate a protein segment.

#### 2.2.3 Building database with alternative splicing events

We use the MATS (Multivariate Analysis of Transcript Splicing) tool (version 4.0.2) ^50^ to identify alternative splicing events from RNA-seq data. MATS reports 5 different splicing events: exon skipping, mutually exclusive exons, alternative 3’ splice sites, alternative 5’ splice sites, and intron retention, in which exon skipping events are the most common form (~30%) ^51^. Here we generate only proteoforms with exon skipping events. An exon splicing event involves three exons: the upstream exon, the downstream exon and the cassette (skipped) exon, and results in two transcript forms: the inclusive form includes the cassette exon, and the exclusive form does not. If a transcript in genome annotation contains the upstream and downstream exons (with or without the cassette exon) of an exon splicing event, then the transcript is matched to the event. We search each exon skipping event against the GENCODE hg38 basic annotation to find all matched transcripts and then generate an inclusive form for each exclusive form transcript matched to the event and *vice versa*. Both the transcript in the hg38 basic annotation and the one generated with the alterative splicing events are added to the customized proteoform database after duplications are removed.

#### 2.2.4 Proteoform identification

The raw files of the DLD-1 MS data were centroided and converted into mzML files using msconvert in ProteoWizard ^52^, and further deconvoluted into msalign files containing monoisotopic masses of fragment and precursor ions using TopFD (version 1.3.1). The deconvoluted spectra were searched against protein databases using TopPIC (version 1.3.1) ^20^. In database search, a shuffled decoy database with the same size was concatenated with the target database. The error tolerances for precursor and fragment masses were set to 15 parts-per-million (ppm). Identifications of proteoform-spectrum-matches (PrSMs) were filtered using a 1% spectrum-level false discovery rate (FDR). Identified proteoforms were treated as the same if their spectra were generated from the same LC-MS feature reported by TopFD or they were from the same *protein* and had similar precursor masses (within an error tolerance of 2.2 Dalton). Using the method, identified PrSMs were grouped into clusters, each of which corresponds to a proteoform. Then the identifications of proteoforms and proteins were filtered using a 1% proteoform-level FDR.

## 3. Results

### 3.1 Comparison of protein reference databases

Three human proteome databases were compared for proteoform identification by database search. The first two databases were generated using the basic and comprehensive annotations of GENCODE (version 28) ^53^ and are referred to as the BASIC (57089 entries) and COMP (97713 entries) databases, respectively. All sequences in BASIC are included in COMP. The third one is referred to as the SWISS (19236 entries) database, which is a subset of BASIC containing only proteoforms matched to entries in the Swiss-Prot human proteome database (20380 entries, March 2019).

We searched the spectra in the DLD-1 top-down MS data against the three proteoform reference databases separately using TopPIC (see Methods). With a 1% spectrum-level FDR, TopPIC identified 3857, 3590, and 3535 PrSMs from the SWISS, BASIC, and COMP databases, respectively (Fig. 1a). We identified more proteoforms and proteins with the SWISS database than the other two. The main reason is that the increase of the database size introduces many decoy identifications in the target-decoy approach and increases the Q-values of identifications in BASIC and COMP. Because of this, some identifications in the BASIC and COMP database search were filtered out. In addition, the BASIC and COMP database searches identified some proteoforms not included in the SWISS database (Fig. 1b).

**Figure 1.**
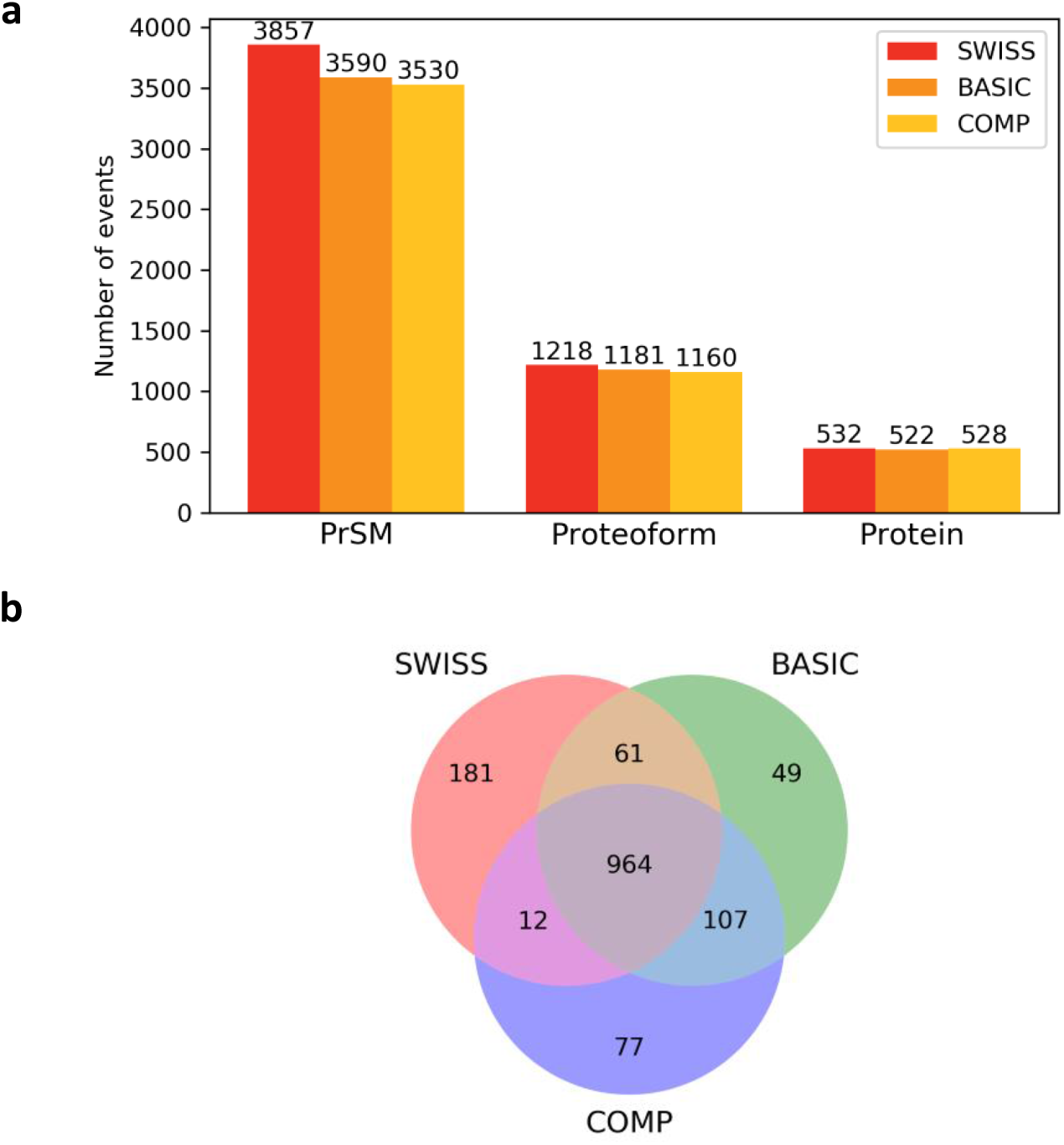
Comparison of the three protein databases SWISS, BASIC, and COMP on proteoform identification using the DLD-1 MS data set. (a) The numbers of identified PrSMs, proteoforms, and proteins. (b) Venn diagram showing the overlaps of the proteoforms identified from the three databases.

### 3.2 Sample specific databases with genomic alterations

The RNA-Seq data of DLD-1 cells were analyzed by the GATK pipeline for short read alignment and SNV calling (see Methods). ANNOVAR ^49^ reported 18133 genomic variants with the basic annotation of GENCODE (version May-06-2018), including 9283 nonsynonymous SNVs, 503 frameshift indels and 152 non-frameshift indels. The nonsynonymous SNVs were mapped to 5420 genes and 14135 transcripts. Of the 14135 transcripts, 36 were not located on the protein-coding region and the others were matched to 5014 and 14099 proteoform sequences in the SWISS and BASIC databases, respectively. The 5014 proteoforms in the SWISS database covered 93% of the 5420 genes with nonsynonymous SNVs. The frameshift indels and non-frameshift indels were also used to generate proteoform sequences with alterations. Using the basic annotation of GENCODE, the indels were mapped to 605 genes and 1553 transcripts, of which 1391 transcripts contained only 1 indel. We used the matched SWISS proteoform sequences to generated a database SWISS-M with 11461 entries: 2633 with homozygous variants only, 6358 with heterozygous variants only, 2470 with both homozygous and heterozygous variants, and used the matched BASIC proteoform sequences to generate a database BASIC-M with 31010 entries: 6840 with homozygous variants only, 17908 with heterozygous variants only, and 6262 with both homozygous and heterozygous variants. Each proteoform sequence in SWISS-M and BASIC-M contained at least one sample-specific genomic variation.

ANNOVAR reported 18682 genomic variants with the comprehensive annotation of GENCODE (9576 nonsynonymous SNVs, 526 frameshift indels, and 154 non-frameshift indels). The nonsynonymous SNVs were mapped to 17983 transcripts of 5549 genes. Similarly, the nonsynonymous SNVs, frameshift, and non-frameshift indels were used to generate a customized database COMP-M with 34276 entries: 7595 with homozygous variants only, 20081 with heterozygous variants only, and 6600 with both homozygous and heterozygous variants.

With a 1% spectrum-level FDR, TopPIC identified 242, 325, and 290 proteoforms from the SWISS-M, BASIC-M, and COMP-M databases, respectively (Fig. 2a). The numbers of identifications were similar for the three databases, and BASIC-M achieved the largest number of identifications. All the identified spectra were matched to database proteoform sequences generated for SNVs, and no identified spectra were matched those for indels. To get the SNV sites covered by the identified proteoforms, we mapped the proteoforms to their corresponding RNA transcript segments and checked whether the transcript segments contain SNV sites. Of the 242 proteoforms identified by SWISS-M, 211 covered 59 SNV sites (some SNV sites were covered by more than one proteoform) and 31 did not cover any SNV sites (Fig. 2c). We further manually inspected the PrSMs covering the 59 SNV sites and found that 42 SNVs were confidently identified. The other 17 SNV sites were not validated because they were covered by proteoforms containing unexpected mass shifts near the SNV sites. Similarly, we manually validated 64 and 61 SNV sites identified from BASIC-M and COMP-M, respectively.

**Figure 2.**
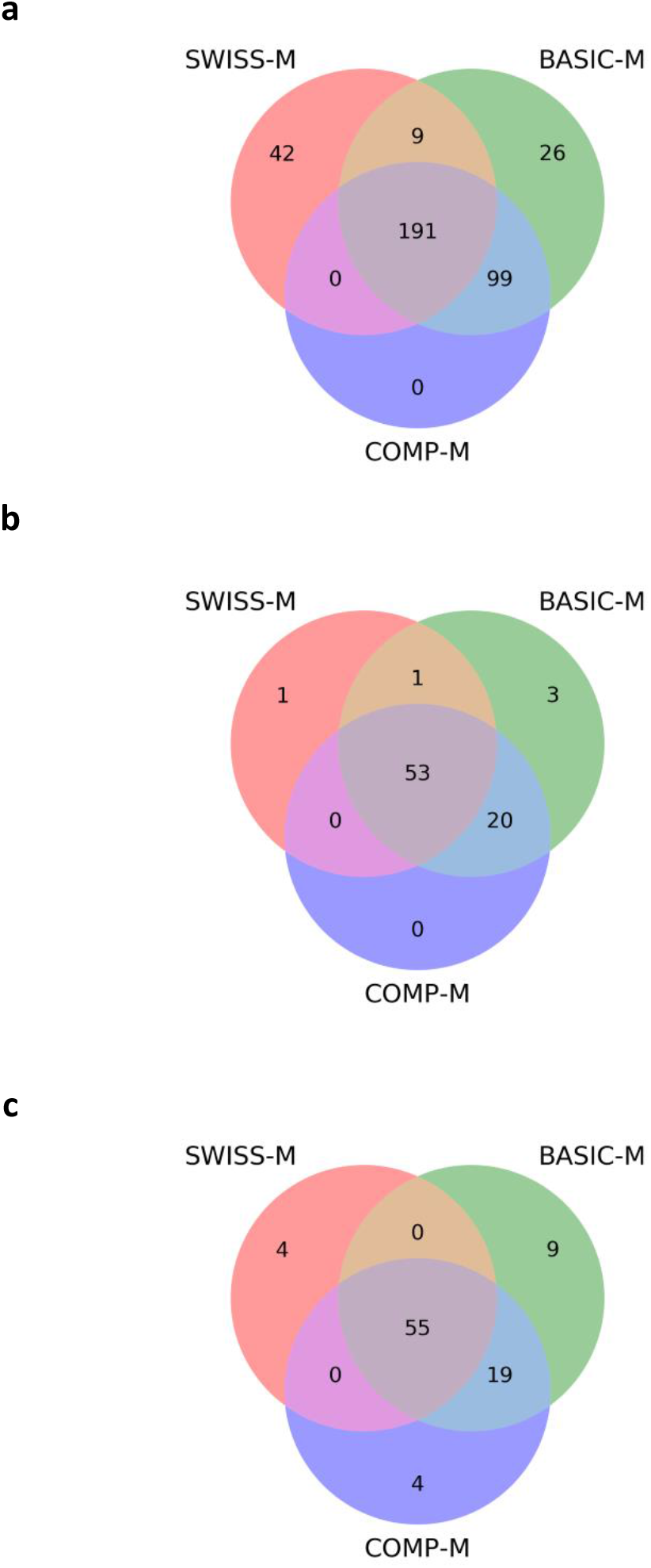
Comparison of the three protein databases SWISS-M, BASIC-M, and COMP-M on proteoform identification using the DLD-1 top-down MS data set: (a) identified proteoforms, (b) identified proteins, and (3) identified mutation sites.

The comparison of the three databases on the numbers of identified proteoforms, proteins, and SNV sites is summarized in Fig. 2. COMP-M and BASIC-M identified similarly numbers of proteoforms and SNVs sites because they have similar database sizes. The reason that SWISS-M identified less SNV sites than the other two databases may be that SWISS-M included less proteoforms in the MS data compared with the other two databases. All the validated SNVs reported by COMP-M were identified by SWISS-M or BASIC-M. In practice, we can combine the search results from the three databases to increase the number of identified SNVs.

### 3.3 Sample specific database with three-step pipeline

We used BASIC and BASIC-M to identify sample specific proteoforms from the DLD-1 MS data with a three-step pipeline (Fig. 3). In the first two steps, we searched the spectra against BASIC and BASIC-M sequentially, in which no mass shifts were allowed. The objective of the two steps was to identify proteoforms without unexpected mass shifts. In the third step, the two databases were concatenated, and DLD-1 spectra were searched against the concatenated database to identify proteoforms in which one unexpected mass shift was allowed in an identified proteoform. Spectra identified in the first step were excluded from the database searches in the next two steps. Similarly, spectra identified in the second step were excluded from the analysis of the third step.

**Figure 3.**
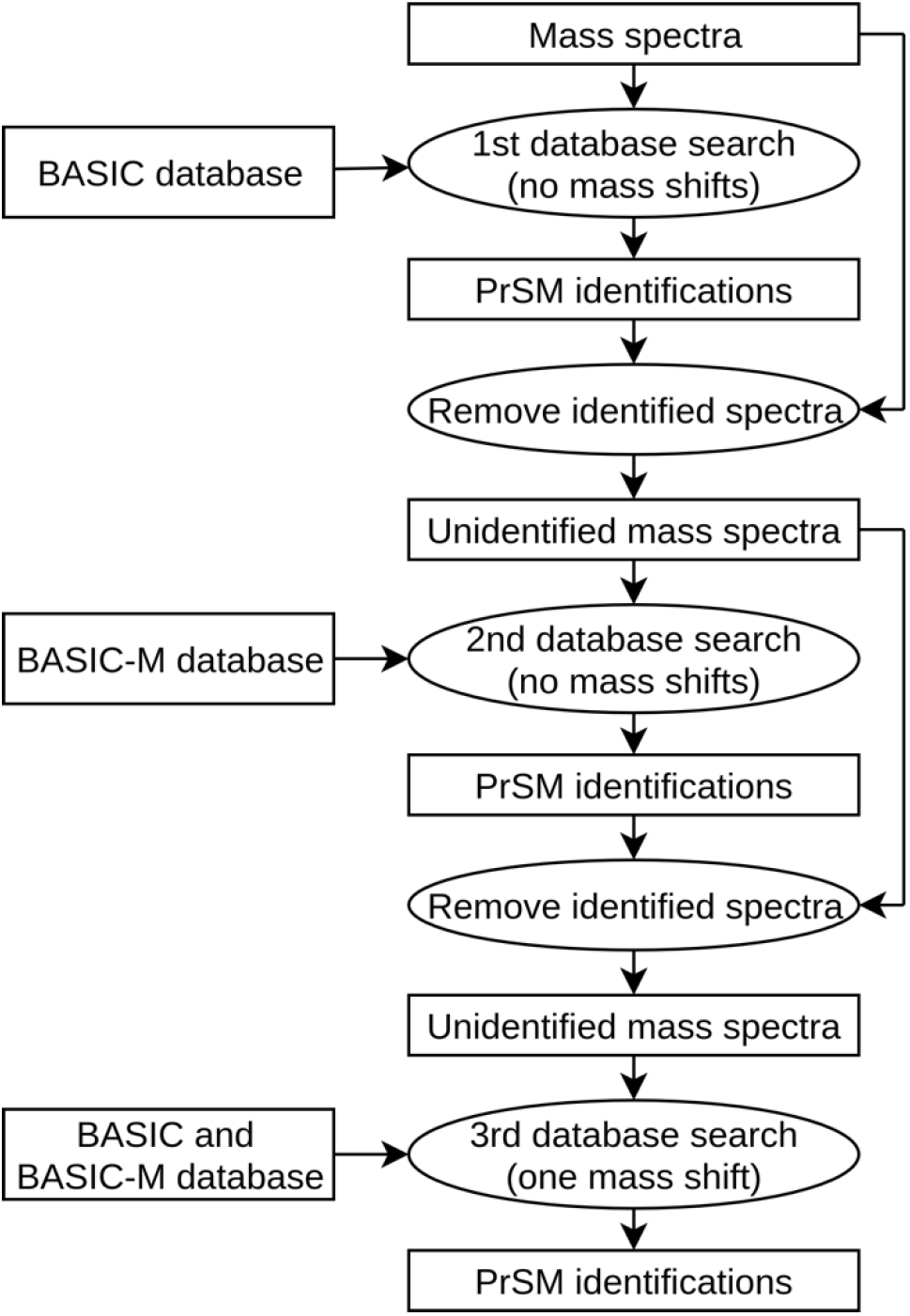
A three-step pipeline for identifying PrSMs from the mass spectra in the DLD-1 data set. In the first two steps, we searched the mass spectra against BASIC and BASIC-M sequentially, in which no mass shifts are allowed. In the third step, the remaining spectra are searched against the concatenated database of BASIC+BASIC-M to identify proteoforms in which one unexpected mass shift is allowed in an identified proteoform.

BASIC-M proteoform identifications covered 27 SNV sites in the second step and 24 SNV sites in the third-step pipeline (Fig. 4). 9 SNV sites were reported in both the second and third steps, so a total of 42 SNV sites were covered by BASIC-M proteoforms. After manual inspection, we removed 7 sites that were not confidently identified. Of the remaining 35 sites, 17 were covered by BASIC-M proteoform identifications only and 18 (10 from the second step and 8 from the third step) were cover by both BASIC and BASIC-M proteoform identifications. The 18 sites covered by both BASIC and BASIC-M proteoforms were all heterozygous.

**Figure 4.**
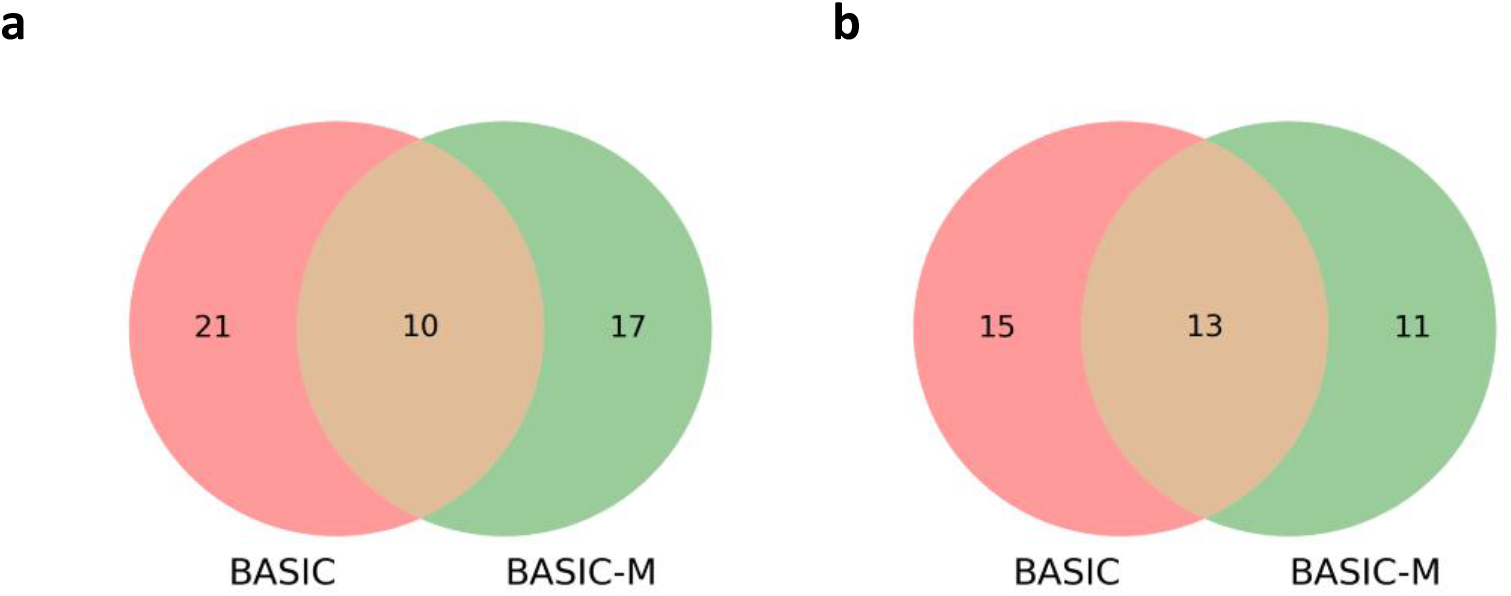
The numbers of identified mutation sites from the DLD-1 data set using a three-step data analysis pipeline. (a) The numbers of mutation sites identified in the first two steps with BASIC and BASIC-M proteoform sequences. (b) the numbers of mutation sites identified in the third step with BASIC and BASIC-M proteoform sequences.

### 3.4 Proteoform identifications with splicing events

MATS ^50^ reported 22774 exon skipping events in the DLD-1 RNA-Seq data. We generated a proteoform databases with 32357 entries, referred to as ES, based on the BASIC annotation of GENCODE. (See Methods.) The top-down mass spectra of DLD-1 cells were searched against the ES database using TopPIC with two steps. First, we searched the MS data against the ES database without unexpected mass shifts and removed identified spectra from the data. Second, we searched the remaining spectra against the ES database in which one unexpected mass shift was allowed in a proteoform.

In the first step, we identified 96 PrSMs and 36 proteoforms from the ES database, of which 24 proteoforms covered 24 splicing events. After manual inspection, we kept 14 inclusive forms and 10 exclusive forms of exon skipping events that were confidently identified. There were no exon skipping events with both inclusive form and exclusive form identifications. In the second step, we identified 219 PrSMs and 94 proteoforms from the database, of which 27 proteoforms covered 28 exon skipping events. Similarly, we manually validated 24 inclusive forms and 4 exclusive forms of exon skipping events that were confidently identified. We found 2 exon-skipping events of which both inclusive forms and exclusive forms were identified. A total of 8 exon-skipping events are covered by two or more proteoforms. In addition, some identified proteoforms cover more than one splicing event.

For each of the inclusive or exclusive form identifications, we computed the RNA expression level of the gene and the percentage of the expressed transcript isoforms containing the exon, called the percent spliced in index (PSI) value, from the DLD-1 RNA-Seq data. The Reads Per Kilobase per Million mapped reads (RPKMs) of genes after logarithm transformation were used as their RNA expression levels. Most of the inclusive form identifications have a high PSI value close to 1.0, and most of the exclusive form identifications have a low PSI value close to 0 (Fig. 5). The splicing events with both inclusive and exclusive forms have a PSI value close to 0.5, showing that the identifications are consistent with the PSI values of the splicing events on the transcript level. In addition, most of the proteoform identifications have a high expression level at the transcript level, demonstrating that top-down MS tends to identify only highly expressed proteoforms.

**Figure 5.**
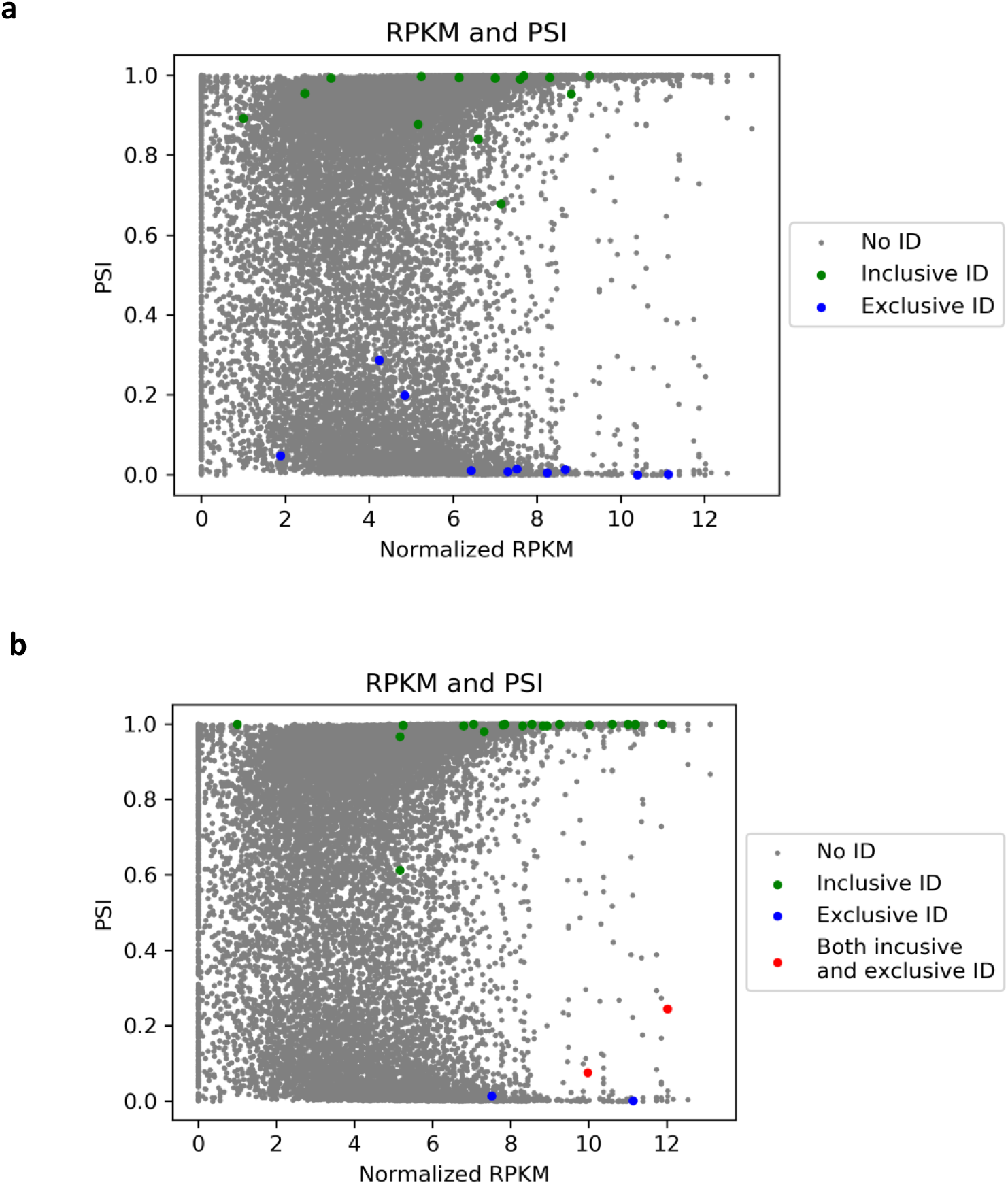
The gene expression levels and PSI values of the splicing events in the DLD-1 data set. The splicing events with proteoform identifications are labeled. (a) Splicing events labeled with proteoform identifications without mass shifts. (b) Splicing events labeled by proteoform identifications with one mass shift.

## 4. Discussion and conclusions

In this study, we proposed a new proteogenomics tool TopPG, which is capable of identifying proteoforms with genomic alterations and alternative splicing events. TopPG builds two types of customized proteoform sequence databases from RNA-Seq data: one for proteoforms with nonsynonymous SNVs, non-frameshift and frameshift indels, the other with exon skipping events. The experiments on the DLD-1 data set demonstrated that databases generated by TopPG facilitated the identification and characterization of sample specific genomic alterations and exon splicing events. In addition, the analysis of exon skipping events showed that their percent spliced in levels were consistent in the transcript and proteoform levels.

Top-down proteogenomics still has many limitations in practice. One limitation is the low proteoform coverage of top-down MS in proteome-level studies. A single shot top-down MS experiment usually identifies only hundreds of proteins, which is a small fraction of genes identified in the transcript level. As a result, only a small number of SNVs and other alterations can be identified by top-down MS.

Another limitation is that some proteoforms with genomic and/or transcriptomic alterations cannot be fully characterized. Manual inspection of the proteoforms identified from the DLD-1 MS data showed that many SNV sites are close to other unknown alterations in identified proteoforms and that the SNV sites cannot be confidently identified because the spectra lack enough fragment ions to characterize them. Similarly, many exon skipping events cannot be confidently identified because the spectra lack enough fragment ions to distinguish between the inclusive and the exclusive forms. Increasing proteoform sequence coverage is essential to identifying sample-specific genomics and transcriptomic alteration.

## 5. Acknowledgements

The research was supported by the National Institute of General Medical Sciences, National Institutes of Health (NIH) through Grants R01GM118470, R01GM125991, and R01AI141625.

## Availability

TopPG is available at http://proteomics.informatics.iupui.edu/software/toppg/.

